# Angiotensin AT1 Receptors Promote Age-Dependent Expansion of Presympathetic Networks in Spontaneously Hypertensive Rats

**DOI:** 10.64898/2026.06.03.729868

**Authors:** Jing-Jing Zhou, Jian-Ying Shao, Shao-Rui Chen, De-Pei Li, Hui-Lin Pan

## Abstract

Heightened sympathetic outflow is a major contributor to the development of hypertension. The hypothalamic paraventricular nucleus (PVN) and the rostral ventrolateral medulla (RVLM) are critical regions for generating and regulating sympathetic activity associated with hypertension. Although presympathetic neural circuitry in the healthy brain is well characterized, it remains unclear whether these pathways undergo alterations in hypertension. Here, we determined presympathetic neural circuits by injecting pseudorabies virus (PRV), a transsynaptic retrograde tracer, into the adrenal gland of spontaneously hypertensive rats (SHR) and normotensive Wistar-Kyoto rats (WKY). Adult SHR exhibited a significantly greater number of PRV-labeled neurons in the PVN and RVLM, but not in the spinal intermediolateral column, compared with WKY. In contrast, the numbers of PRV-labeled neurons in the PVN and RVLM were comparable between young, prehypertensive SHR and age-matched WKY. Remarkably, long-term treatment with losartan—a brain-penetrant angiotensin II AT1 receptor antagonist— initiated in young, prehypertensive SHR blunted the age-dependent hypertension development and reversed the increase in neuronal labeling in both the PVN and RVLM. However, losartan treatment had no effects in WKY. Additionally, electrophysiological recordings showed an elevated frequency of miniature excitatory postsynaptic currents in PVN presympathetic neurons of SHR, which was also normalized by long-term losartan treatment. These findings reveal an age-dependent expansion of presympathetic neuronal connectivity from the hypothalamus and brainstem to the adrenal gland during hypertension development in SHR. Enhanced AT1 receptor activity contributes to hypertension by increasing active glutamatergic synaptic input and promoting the recruitment of additional presympathetic neurons in the hypothalamus and brainstem.

**Key Points:** 1. The numbers of neurons labeled by PRV injected into the adrenal gland are increased in the PVN and RVLM, but not in the spinal cord IML, in adult SHR compared to normotensive WKY. 2. The numbers of neurons in the PVN, RVLM, and spinal cord labeled by PRV injected into the adrenal gland are comparable in young, prehypertensive SHR and age-matched WKY. 3. Losartan treatment, initiated at a young age, blunts the hypertension development and reverses the increased numbers of PRV-labeled neurons in the PVN and RVLM of adult SHR but has no such effects in WKY. 4. The active glutamatergic synapses in PVN presympathetic neurons are elevated in adult SHR, and this elevation is reversed by long-term losartan treatment.

## Introduction

Hypertension is a highly prevalent condition and a major risk factor for cardiovascular disease, stroke, and renal failure. Although the underlying mechanisms of essential hypertension are incompletely understood, the condition likely arises from a complex interplay between genetic and environmental factors (1-4). Increased sympathetic outflow from the brain can cause widespread vasoconstriction, leading to hypertension (5-7). The sympathetic nervous system operates across multiple levels—from central to peripheral—including autonomic centers in the brain (e.g., hypothalamus and brainstem), the intermediolateral column (IML) of the spinal cord, sympathetic ganglia, and postganglionic neurons. However, the precise sites and mechanisms responsible for the persistent increase in sympathetic vasomotor activity in hypertension remain unclear.

The hypothalamic paraventricular nucleus (PVN) is a highly organized structure that coordinates neuroendocrine and autonomic functions involved in stress responses, cardiovascular regulation, thermoregulation, and other physiological processes (7-9). PVN presympathetic neurons project directly to vasopressor neurons in the rostral ventrolateral medulla (RVLM) and to preganglionic neurons in the IML of the spinal cord, thereby regulating sympathetic outflow to the adrenal gland, blood vessels, heart, and kidneys (10-12). Accordingly, the PVN is often referred to as an “autonomic master controller” and represents a major source of excessive sympathetic drive in spontaneously hypertensive rats (SHR) (7, 13, 14), a genetic model of essential hypertension. SHR are normotensive at a young age but develop progressive, age-dependent hypertension (15-17). Electrical lesioning or pharmacological inhibition of the PVN reduces sympathetic nerve discharges and blood pressure in SHR (13, 14, 18). Similarly, neuronal activity in the RVLM is elevated and contributes to the long-term maintenance of hypertension in SHR (19, 20). Although sympathetic-related neuronal circuitry has been mapped in the healthy brain (21-23), it remains unclear whether this circuitry is altered during the development of hypertension in SHR.

An enhanced renin-angiotensin system (RAS) plays a key role in the pathogenesis of hypertension by increasing sympathetic outflow from central autonomic centers (24-28). Sustained blockade of AT1 receptors with losartan, initiated early in life, largely prevents the development of hypertension in SHR (17, 29). In addition to the circulating, kidney-derived RAS, a local brain RAS—particularly within the PVN and RVLM—also contributes to hypertension in SHR (3, 20, 26, 30). The age-dependent rise in blood pressure in SHR is closely linked to potentiated angiotensin II type 1 (AT1) receptor activity in the PVN. For example, AT1 receptor expression in the PVN increases in parallel with the progressive elevation of arterial blood pressure in SHR (3). Systemically administered losartan can cross the blood-brain barrier to block AT1 receptors in the brain (31, 32). However, whether long-term AT1 receptor antagonism affects the brain’s presympathetic neural networks in SHR is largely unknown.

In this study, we used pseudorabies virus (PRV)-152 expressing enhanced green fluorescent protein (GFP), a powerful tool for transsynaptic neuronal labeling (21, 33-35), to investigate potential changes in presympathetic neural circuits in SHR. Our study reveals that the number of presympathetic neurons projecting to the adrenal gland is markedly increased in both the PVN and RVLM of SHR in an age-dependent manner. Strikingly, long-term blockade of AT1 receptors with losartan, initiated early in life, reverses this expansion of presympathetic neuronal connectivity by reducing the heightened excitatory synaptic activity of these neurons. These findings suggest that enhanced AT1 receptor activity drives the expansion of presympathetic neuronal networks in the brain, contributing to sustained sympathetic overactivity and the development of neurogenic hypertension.

## Materials and Methods

### Animal model

The experimental protocol (#919-RN03) was approved by the Institutional Animal Care and Use Committee of The University of Texas MD Anderson Cancer Center and conformed to the National Institutes of Health guidelines on the ethical use of animals. Young (4 weeks old) and adult (12 weeks old) male SHR and age-matched male Wistar-Kyoto rats (WKY) were used in the study. SHR and WKY were purchased from Harlan Laboratories (Envigo) and housed (3 rats per cage) on a 12-hour light/dark cycle with free access to food and water at the animal facility. Tail-cuff method was used to confirm hypertension in adult SHR.

Losartan was purchased from MedChem Express (#HY-17512, Monmouth Junction, NJ) and given to rats at a dose of 50 mg/kg per day via drinking water. Rats receiving long-term losartan and vehicle treatment were housed individually. All efforts were made to minimize animal suffering and the number of animals used.

### PRV-152 injection

PRV-152 (3 × 10^8^ plaque forming units/mL) was obtained from the Viral Vector Core Facility at Princeton University (Princeton, NJ). PRV-152 carried cytomegalovirus-enhanced green fluorescent protein (GFP) reporter gene cassette, which was inserted into the gG locus of viral genome (36, 37). PRV-152 stock was aliquoted in 5.0 µL volumes and stored at –80°C. Rats were anesthetized with 3% isoflurane, and the left adrenal gland was exposed via a left flank incision and gently separated from the surrounding viscera and fat. Two separate 1.0 µL injections of PRV-152 (total 2.0 µl) were administered into the left adrenal gland (22, 23, 33). All injections were performed using a 30-gauge steel needle attached to a Hamilton syringe (10.0 µL). The needle was held with a micromanipulator, and the viral suspension was slowly injected over a period of 5 min. As soon as the needle was withdrawn from the adrenal gland, a drop of liquid antiseptic plastic sealant (#370-0005-001, Data Sciences International) was applied to prevent viral leakage. The surgical wound was closed in layers. Animals were then given prophylactic enrofloxacin (0.5 mg/100 g, daily for 3 days) and subcutaneous buprenorphine-ER (0.1 mg/100 g, once every 48 hours for a total of two doses). Animals receiving PRV-152 injections were housed in a designated biohazard housing room within the animal facility.

### Tissue processing and immunofluorescence labeling

Rats injected with PRV-152 were sacrificed for tissue fixation on day 6 post PRV-152 injection. Under deep anesthesia with sodium pentobarbital (60 mg/kg, i.p.), rats were perfused intracardially with 0.9% saline followed by 4% paraformaldehyde and then 10% sucrose in 0.1 M phosphate-buffered saline (PBS). The brain and spinal cord were carefully removed, post-fixed in 4% paraformaldehyde at 23°C for 1 hour, and then cryoprotected in 30% sucrose in 0.1 M PBS at 4°C for 48 hours. Brain sections were cut coronally at 30 μm, and spinal cord sections were cut sagittally at 30 μm. The tissue sections were incubated with a rabbit anti-Neuronal Nuclei (NeuN) antibody (1:300; #ab177487, Abcam) in 0.1 M PBS containing 3% goat serum at 4°C for 24 hours. On the following day, the sections were rinsed and incubated with goat anti-rabbit-Alexa Fluor-594 (1:500; #111-585-003, Jackson ImmunoResearch Laboratories) in 0.1 M PBS containing 3% goat serum at 23°C for 1 hour. The sections were then rinsed in 0.1 M PBS for 10 min, three times, and finally mounted on glass slides using mounting medium (#ab104139, Abcam).

### Image processing and cell counting

Preganglionic neurons innervating the adrenal gland are located in the spinal cord IML (0.8–1.0 mm lateral to the spinal cord midline) at T5–T9 and are predominantly concentrated at T7–T10 (23). In transverse sections, the IML at T7–T10 appears as a triangular horn plane measuring approximately 200 µm mediolaterally and 120 µm dorsoventrally (38). The spinal cord was cut sagittally, and 6 sagittal sections covering the IML from T7 to T10 were used for cell counting.

The RVLM and PVN were identified according to the rat brain atlas (38). The RVLM extends 1.5 mm caudally from the caudal pole of the facial nucleus (–11.3 mm to –12.8 mm posterior to bregma) and is bounded by the apex of the nucleus ambiguous, the ventral border of the spinal trigeminal tract, and the lateral border of the pyramidal tract (38). A total of 48 coronal RVLM sections were cut per animal, spanning caudal to rostral. Every 6th section was selected, yielding 8 sections per animal for quantification. The PVN is defined as 1.4–2.2 mm caudal to bregma, 0.3–0.5 mm lateral to the midline, and 7.7–8.0 mm below the dorsal surface (38). Because presympathetic neurons in the PVN are distributed at mid-rostrocaudal to caudal levels (12), a total of 12 coronal PVN sections were cut across this range. Every 4th section was selected, resulting in 3 sections per animal for cell counting.

PRV-labeled neurons were visualized and imaged using a confocal microscope (model LSM510, Carl Zeiss Microscopy). Fluorescent signal quantification was performed using a hybrid cell-counting module (Keyence Corporation of America), which analyzed pixel brightness to quantify fluorescent objects. The pixels in non-cellular regions were measured to establish a fluorescence-intensity threshold used to distinguish labeled cells from background. Cells exceeding this threshold and with a diameter larger than 6.0 µm were counted (39). Consistent with previous findings, PRV-labeled neurons were predominantly located on the side ipsilateral to the injection site (22, 40). Therefore, only ipsilateral PRV-labeled neurons were quantified for group comparisons. For each animal, the number of NeuN-positive and PRV-labeled neurons within each section from the same region was summed to obtain a total count, and these totals were averaged across animals within each group. Cell counting was performed by an investigator blinded to both group and treatment.

### Retrograde labeling of spinally projecting PVN neurons

We used a fluorescent microsphere tracer (FluoSpheres with diameter 0.04 μm, Invitrogen) to retrogradely label spinally projecting PVN presympathetic neurons as described previously (26, 41). Briefly, rats were anesthetized with 3–4% isoflurane, and dorsal laminectomy was performed to expose the spinal cord at the upper thoracic level. The fluorescent microsphere suspension was pressure-ejected (Nanoject II; Drummond Scientific Company) into the IML through a glass pipette (tip diameter: 20–30 μm) in 3 separated 50-nl injections on each side, monitored under a surgical microscope. Animals were then treated prophylactically with enrofloxacin (5 mg/kg, daily for 3 days) to prevent infection and with subcutaneous buprenorphine-SR (0.1 mg/100 g, once every 48 hours for a total of two times) to minimize pain. The animals were allowed to recover for 5 days to permit retrograde transport of the FluoSpheres to the PVN.

### Hypothalamus slice preparation and whole-cell recordings

Rats were anesthetized with 5% isoflurane and then rapidly decapitated. The brain was removed and placed in an ice-cold artificial cerebrospinal fluid containing 124.0 NaCl, 3.0 KCl, 1.3 MgSO_4_, 2.4 CaCl_2_, 1.4 NaH_2_PO_4_, 10.0 glucose, and 26.0 NaHCO_3_, which was saturated with a mixture of 95% O_2_ and 5% CO_2_. A tissue block containing the PVN was trimmed from the brain tissue and glued onto the stage of a vibrating microtome (VT1000s, Leica). The tissue block was cut coronally at 300 μm thickness, and the brain slices were incubated in the artificial cerebrospinal fluid continuously gassed with a mixture of 95% O_2_ and 5% CO_2_ at 34°C for at least 1 hour before electrophysiological recordings.

The slices were transferred into a recording chamber and continuously perfused with the artificial cerebrospinal fluid at a speed of 3 ml/min at 34°C. Because spinally projecting PVN presympathetic neurons are primarily concentrated in the mediocellular region of the PVN (42), we selected fluorescence-labeled neurons in this region for electrophysiological recordings. Labeled PVN neurons were identified using an upright microscope equipped with epifluorescence illumination and differential interference contrast optics (BX50 WI; Olympus Optical). All electrophysiological recordings were performed using the whole-cell voltage-clamp configuration. Recording electrodes were pulled from borosilicate capillaries (1.2 mm outer diameter, 0.68 mm inner diameter; World Precision Instruments) using a micropipette puller (P-97, Sutter Instruments). The electrode resistance was 4.0–5.0 MΩ when filled with an internal solution containing (in mM): 140.0 K gluconate, 2.0 MgCl_2_, 0.1 CaCl_2_, 10.0 HEPES, 1.1 EGTA, 0.3 Na_2_-GTP, 2.0 Na_2_-ATP and 10 lidocaine N-ethyl bromide (QX-314) adjusted to pH 7.25 with 1 M KOH and osmolality of 270–290 mOsm. Miniature excitatory postsynaptic currents (mEPSCs) were recorded at a holding potential of –60.0 mV in the presence of 1.0 μM tetrodotoxin (#HB1035, Hello Bio) (26, 41). Signals were amplified with a Multiclamp 700B amplifier, filtered at 1.0–2.0 kHz, and digitized at 20.0 kHz using DigiData 1550B (Molecular Devices).

### Data and statistical analyses

All data are presented as means ± S.D. The group sample sizes were comparable to those commonly used in the field (14, 21, 26, 27, 40, 43). The animals were assigned (1:1 allocation) to the control and treatment groups as they became available, without the use of specific randomization methods. Investigators were blinded to the animals’ genotype and treatment during electrophysiological recordings and cell counting. Representative microscopic images and electrophysiological recording traces were chosen to best reflect the mean data. Male rats were used in this study, because SHR develop age-dependent hypertension and increased AT1 receptor expression in the PVN that occur independently of sex (3). For the electrophysiology experiments, recordings were obtained from a single neuron in each brain slice, with at least three rats included in each experimental group. Cell capacitance and series resistance were continuously monitored during recordings; cells were excluded if series resistance changed by more than 20%. The amplitude and frequency of mEPSCs were analyzed using a peak detection algorithm with a threshold set above the noise level (MiniAnalysis, Synaptosoft). Cumulative probability distributions of mEPSC amplitude and inter-event intervals were compared using the Kolmogorov–Smirnov test to assess distributional differences. Data normality was evaluated with the Shapiro-Wilk test. For statistical comparisons, two-tailed Student’s t tests were used for two-group analyses. Differences among three or more groups were assessed using two-way ANOVA followed by Tukey’s post hoc test. Repeated one-way ANOVA with Dunnett’s post hoc test was used to analyze differences across time points within a group. All statistical analyses were performed using GraphPad Prism (Version 10.3). A *P* value < 0.05 was considered statistically significant.

## Results

### The number of spinal IML neurons labeled by PRV injected into the adrenal gland is comparable between adult SHR and WKY

PRV-152 is an attenuated PRV-Bartha strain that expresses GFP and is widely used as a transsynaptic retrograde tracer (33, 34). We selected the adrenal medulla for PRV injection because it receives dense, direct innervation from preganglionic sympathetic neurons in the spinal IML (23). Furthermore, the adrenal sympathetic nerve activity is markedly increased in SHR (44), and adrenal demedullation largely reduces plasma catecholamine levels and attenuates the development of hypertension in SHR (45-47). Adult SHR (12 weeks old) exhibit persistent hypertension compared with normotensive WKY. To map presympathetic neuronal connections to the adrenal gland, PRV-152 was injected into the left adrenal medulla of adult SHR and WKY (n = 6 rats per group). Rats were sacrificed 6 days after injection, a time point shown to produce maximal neuronal labeling (40). Both the spinal cord and brain were collected for neuronal immunostaining with NeuN and neuronal cell counting. The adrenal gland receives sympathetic input from preganglionic neurons in the IML at spinal cord levels T3−L2, predominantly on the ipsilateral side (48, 49). Also, presympathetic neurons in the RVLM and PVN primarily project ipsilaterally to sympathetic preganglionic neurons in the IML, which in turn innervates the adrenal medulla (22, 50). Therefore, quantification of labeled neurons was focused on the IML, RVLM, and PVN ipsilateral to the adrenal gland that received the PRV-152 injection.

We first determined whether the number of PRV-labeled IML neurons innervating the adrenal medulla differs between SHR and WKY. IML preganglionic neurons are mostly concentrated at T7–T10 levels of the spinal cord (23). To quantify these neurons, PRV-labeled IML neurons were counted on sagittal spinal cord sections, because the number of labeled IML neurons was low in transverse sections in the pilot study (**Fig. 1A**). All PRV-labeled neurons in the IML were immunoreactive to NeuN, a pan-neuronal marker (**Fig. 1B,C**). The distribution pattern and total number of PRV-labeled neurons in the spinal IML were comparable between SHR and WKY rats (**Fig. 1B,C**). These data suggest that the connectivity between IML preganglionic neurons and the adrenal gland is unaltered in SHR.

**Fig. 1.**
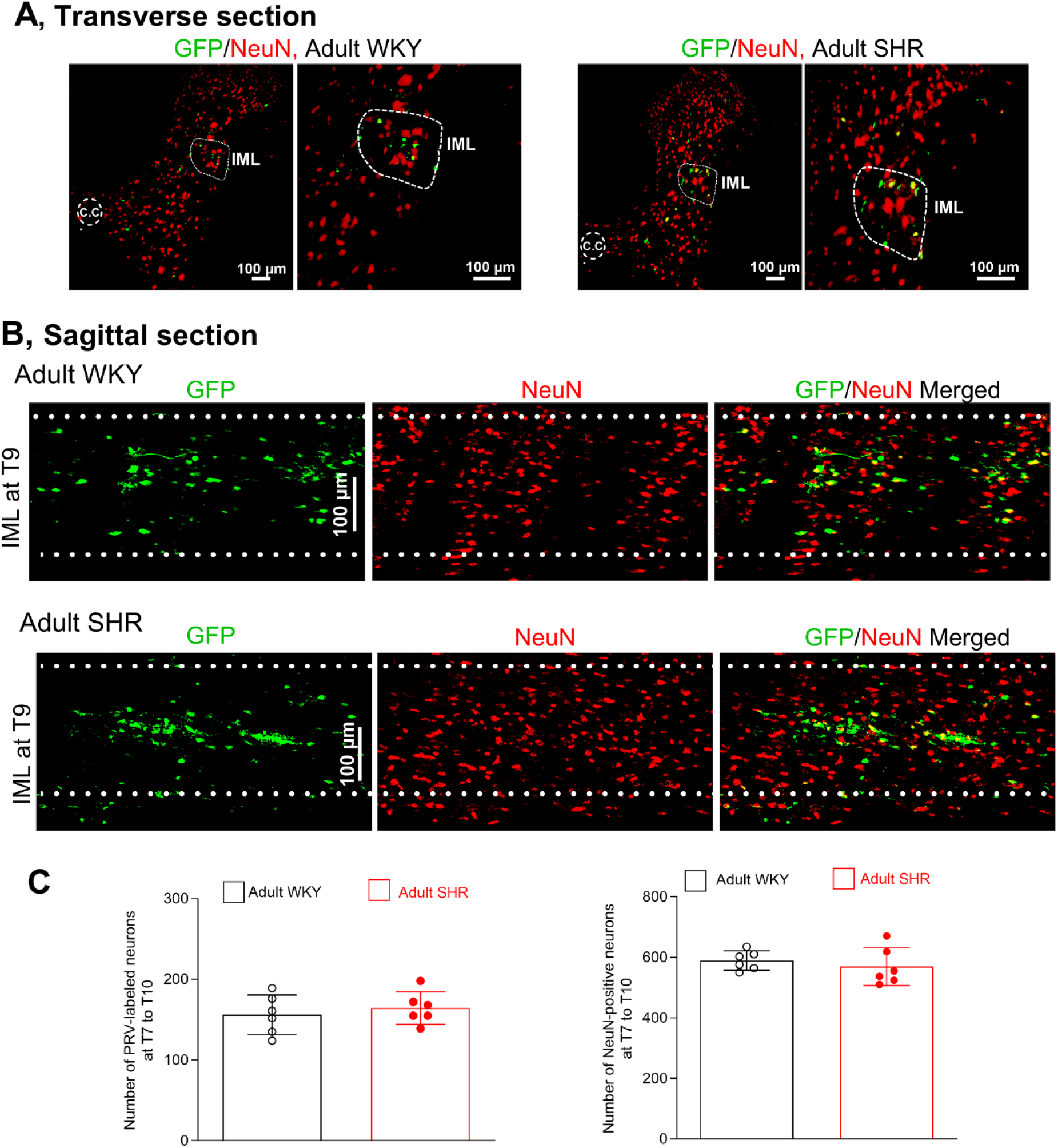
The number of preganglionic neurons innervating the adrenal gland is unaltered in the spinal IML of adult SHR. **A**, Representative low- and high-magnification images of transverse sections at the T9 level show dual labeled GFP and NeuN neurons in the spinal cord IML of adult WKY and SHR with PRV-152 injected into the left adrenal gland. CC, central canal. Scale bar, 100 µm. **B**, Representative confocal images of sagittal spinal cord sections at the T9 level show preganglionic neurons (dual-labeled by GFP and NeuN) in the spinal IML of adult WKY and SHR with PRV-152 injected into the left adrenal gland. Scale bar, 100 µm. **C**, Summary data show the number of preganglionic neurons (dual-labeled by GFP and NeuN) and NeuN-immunoreactive neurons in the spinal cord IML from T7 to T10 levels in adult WKY and SHR (n = 6 rats per group; two-tailed Student’s *t* test).

### The number of RVLM neurons labeled by PRV injected into the adrenal gland is increased in adult SHR

Presympathetic neurons in the RVLM form direct connections with spinal IML preganglionic neurons (10, 22). We next quantified PRV-labeled neurons in the RVLM of adult SHR and WKY. The distribution pattern of PRV-labeled neurons within the RVLM was similar between the two groups (**Fig. 2A,B**). Also, the total number of NeuN-positive neurons in the RVLM did not differ between the two groups (**Fig. 2A-C**). However, the number of PRV-labeled neurons in the RVLM was significantly greater in SHR than in WKY (n = 6 rats per group, *P* < 0.001, *t*_(10)_ = 13.96; **Fig. 2A-C**). These results suggest that presympathetic neuronal connectivity from the RVLM to the adrenal gland is enhanced in adult SHR.

**Fig. 2.**
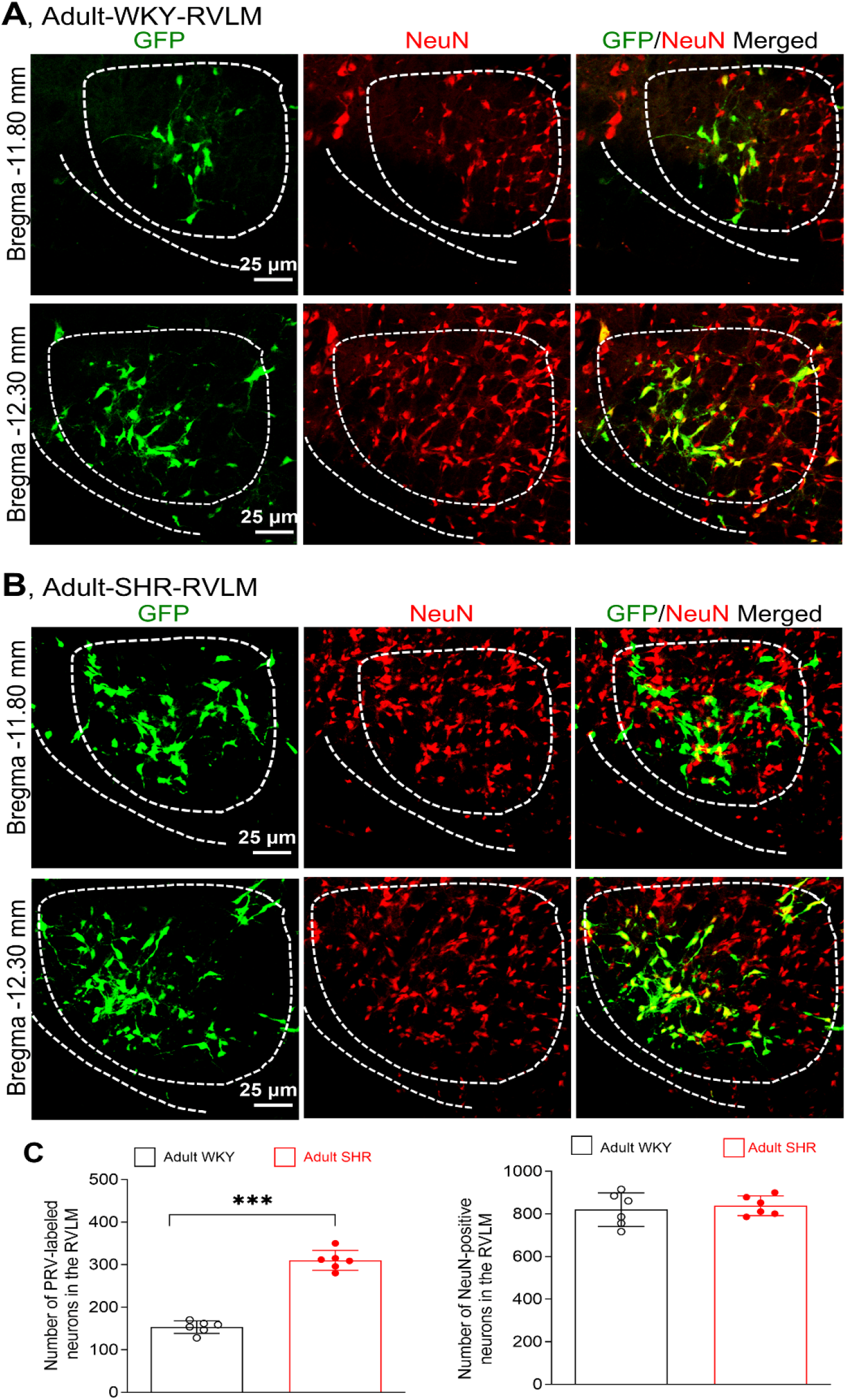
The number of presympathetic neurons innervating the adrenal gland is increased in the RVLM of adult SHR. **A** and **B**, Representative confocal images show presympathetic neurons (dual-labeled by GFP and NeuN) in the RVLM of adult WKY (**A**) and SHR (**B**) with PRV-152 injected into the left adrenal gland. Scale bar, 25 µm. **C**, Quantification of the number of presympathetic neurons (dual-labeled by GFP and NeuN) and NeuN-immunoreactive neurons in the RVLM of adult SHR and WKY (n = 6 rats per group; \*\*\**P* < 0.001, two-tailed Student’s *t* test).

### The number of PVN neurons labeled by PRV injected into the adrenal gland is increased in adult SHR

Presympathetic neurons in the PVN send direct projections to the spinal IML and the RVLM (11). We therefore determined whether the number of PVN presympathetic neurons projecting to the adrenal gland differs between adult SHR and WKY. PRV-labeled neurons were distributed primarily at middle to caudal levels of the PVN in both groups (**Fig. 3A,B**). The overall distribution pattern was similar between SHR and WKY. Furthermore, the total number of NeuN-positive neurons in the PVN did not differ between the two groups (**Fig. 3A-C**). However, the number of PRV-labeled neurons in the PVN was substantially higher in SHR than in WKY (n = 6 rats in each group, *P* < 0.001, *t*(_10)_ = 9.38; **Fig. 3A-C**). These findings suggest that PVN presympathetic neuronal connectivity to the adrenal gland is augmented in adult SHR.

**Fig. 3.**
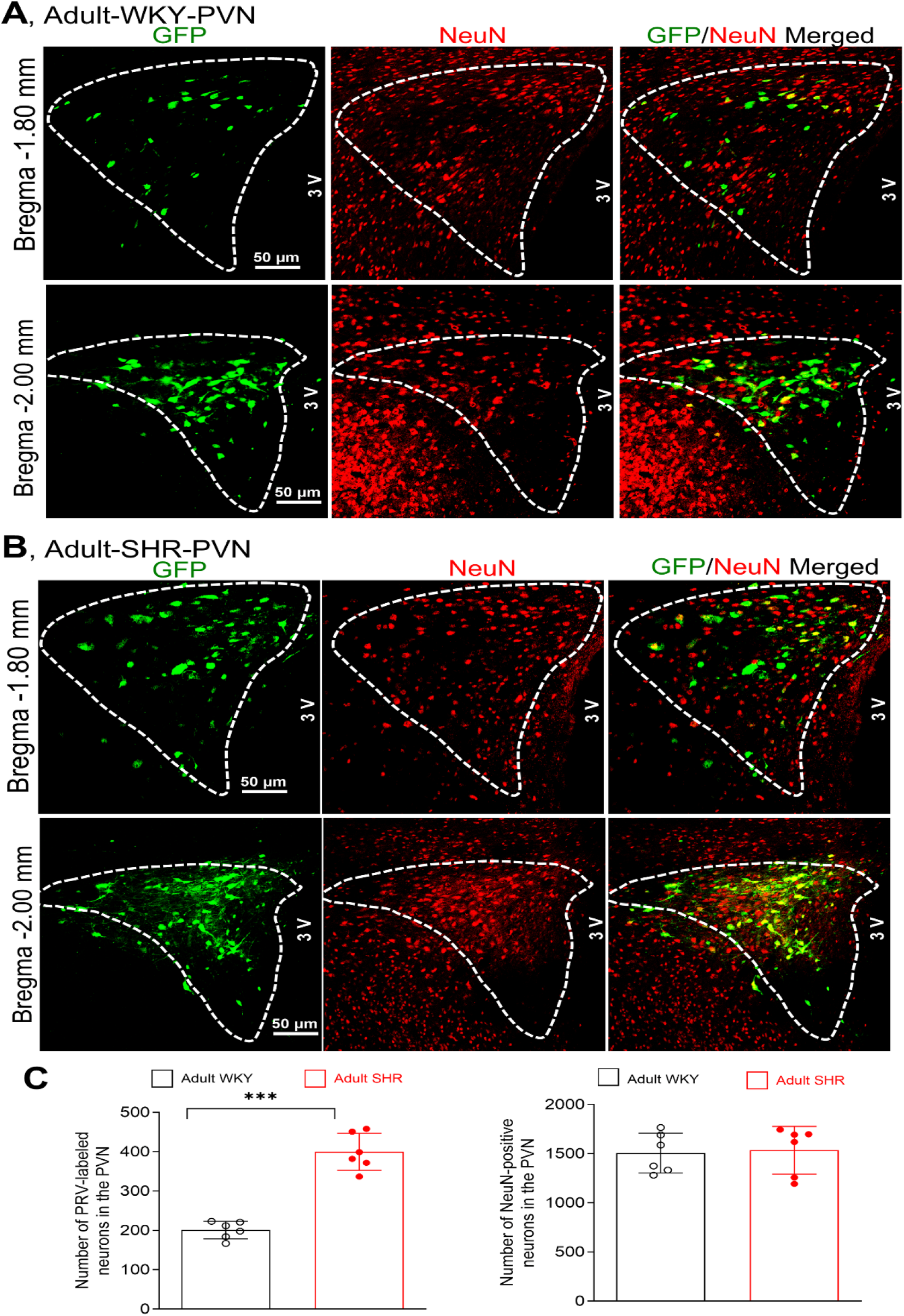
The number of presympathetic neurons innervating the adrenal gland is increased in the PVN of adult SHR. **A** and **B**, Representative confocal images show presympathetic neurons (dual-labeled by GFP and NeuN) in the PVN of adult WKY (**A**) and SHR (**B**) with PRV-152 injected into the left adrenal gland. Scale bar, 50 µm. **C**, Summary data show the number of presympathetic neurons (dual-labeled by GFP and NeuN) and NeuN-immunoreactive neurons in the PVN of adult SHR and WKY (n = 6 rats per group; \*\*\**P* < 0.001, two-tailed Student’s test).

### The numbers of PVN and RVLM neurons labeled by PRV injected into the adrenal gland are comparable in young, prehypertensive SHR and WKY

In SHR, arterial blood pressure remains within the normotensive range until approximately 4 weeks of age, begins to rise progressively around 5–6 weeks, and reaches a sustained hypertensive level by 12 weeks of age (51). Because the numbers of presympathetic neurons in the RVLM and PVN are significantly increased in adult SHR, we next determined whether this increased presympathetic neuronal connectivity is already present in young, prehypertensive SHR. PRV-152 was injected into the left adrenal gland of 4-week-old SHR and WKY, and rats were euthanized 6 days after injection. The brain and spinal cord tissues were collected for PRV-labeled cell counting. Injection of PRV-152 into the adrenal gland resulted in similar numbers of labeled neurons in the IML, RVLM, and PVN between young SHR and WKY (**Fig. 4; Suppl Fig. S1**). The distribution pattern of PRV-labeled neurons in these regions was also comparable between the two groups (**Fig. 4; Suppl Fig. S1**). These data indicate that the expanded brain presympathetic connectivity to the adrenal gland in adult SHR is not congenital but instead develops over time, coinciding with the rise in blood pressure.

**Fig. 4.**
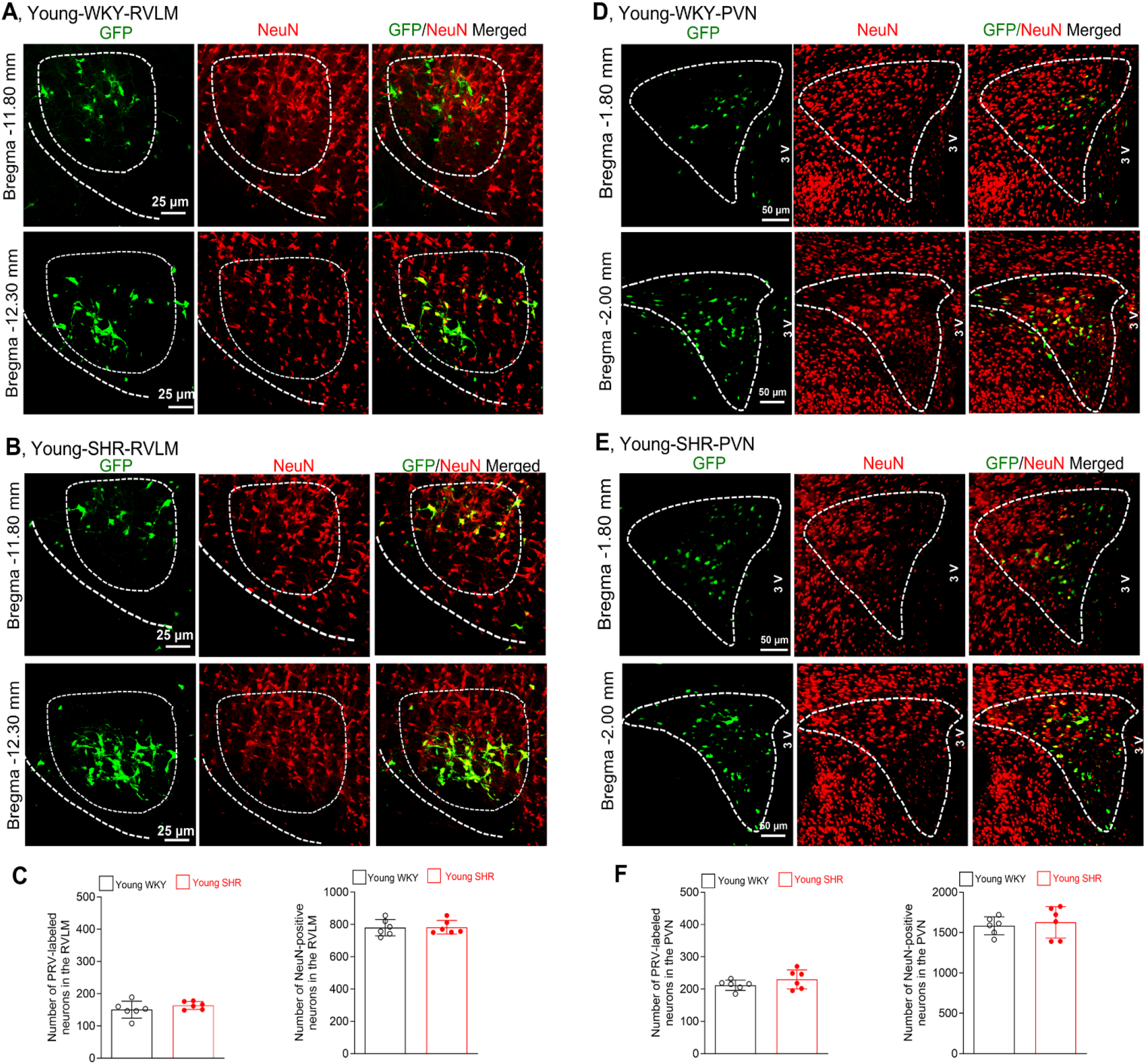
The numbers of presympathetic neurons innervating the adrenal gland are unaltered in the RVLM and PVN of young, prehypertensive SHR. **A–C**, Representative confocal images (**A** and **B**) and quantification (**C**) show the number of presympathetic neurons (dual-labeled by GFP and NeuN) and NeuN-immunoreactive neurons in the RVLM of young WKY and SHR with PRV-152 injected into the left adrenal gland (n = 6 rats per group). Scale bar, 25 µm. **D–F**, Representative confocal images (**D** and **E**) and summary data (**F**) show the number of presympathetic neurons (dual-labeled by GFP and NeuN) and NeuN-immunoreactive neurons in the PVN of young WKY and SHR with PRV-152 injected into the left adrenal gland (n = 6 rats per group; two-tailed Student’s *t* test). Scale bar, 50 µm.

### Long-term treatment with losartan diminishes the increase in brain presympathetic neurons labeled by PRV injected into the adrenal gland in SHR

Subsequently, we sought to define the mechanisms underlying the expansion of presympathetic neural networks in adult SHR. Potentiated AT1 receptor expression and activity in the PVN and RVLM contribute to elevated sympathetic outflow in SHR (3, 26, 52). Importantly, long-term administration of losartan, a specific angiotensin II AT1 receptor antagonist, in young, prehypertensive SHR largely blocks the development of hypertension (17, 29). Because losartan readily crosses the blood-brain barrier (31, 32), we investigated whether prolonged AT1 receptor blockade influences the expansion of presympathetic neuronal connectivity in the brain of adult SHR. SHR and WKY were treated with losartan (50 mg/kg per day in drinking water) (16, 53) or vehicle from 4 to 12 weeks of age. In vehicle-treated SHR, systolic arterial blood pressure, measured using the tail-cuff method, increased progressively from 5 weeks of age and reached a stable hypertensive level by 12 weeks (n = 6 rats, *F*_(5,70)_ = 89.0, *P* < 0.001; **Fig. 5A**). As expected, losartan treatment for 8 weeks blunted this age-dependent rise in arterial blood pressure in SHR (n = 6 rats, *F*_(3,20)_ = 125.98, *P* < 0.001; **Fig. 5A**). In contrast, long-term treatment with losartan had no evident effect on arterial blood pressure in WKY (n = 6 rats per group, **Fig. 5A**).

**Fig. 5.**
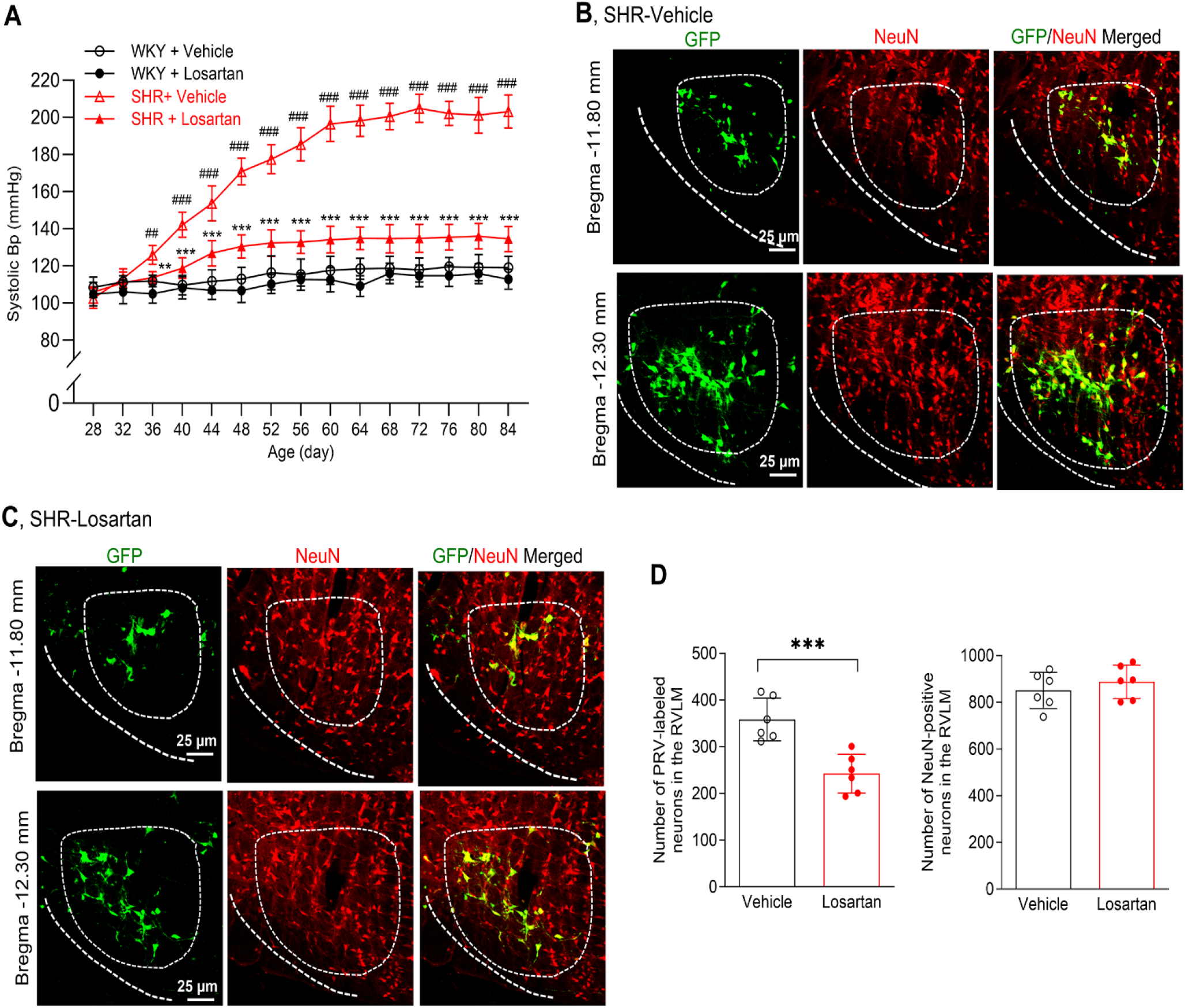
Long-term treatment with losartan blunts age-dependent hypertension development and the increased number of presympathetic neurons innervating the adrenal gland in the RVLM of SHR. **A**, Time course of systolic arterial blood pressure (Bp) in WKY and SHR treated with vehicle or losartan (50 mg/kg per day), from 4 to 12 weeks of age. Two-way ANOVA with Tukey *post hoc* test was used for comparisons among the four groups; repeated ANOVA with Dunnett’s *post hoc* test was used for comparison within the group. \*\**P* < 0.01 and \*\*\**P* < 0.001 compared with the SHR+vehicle group at the same time point; ^*##*^*P* < 0.01 and ^*###*^*P* < 0.001 compared with the Bp value at day 28 in the SHR+vehicle group. **B–D**, Representative confocal images (**B** and **C**) and summary data (**D**) show the number of presympathetic neurons (dual-labeled by GFP and NeuN) and NeuN-immunoreactive neurons in the RVLM of vehicle- or losartan-treated SHR from 4 to 12 weeks of age followed by PRV injection into the left adrenal gland (n = 6 rats per group; \*\*\**P* < 0.001, two-tailed Student’s *t* test). Scale bar, 25 µm.

PRV-152 was injected into the left adrenal gland of adult SHR and WKY after discontinuation of losartan or vehicle treatment. The brain and spinal cord tissues were obtained 6 days after PRV-152 injection for counting of labeled neurons. The distribution patterns of PRV-labeled neurons in the spinal IML, RVLM, and PVN were similar between vehicle-treated and losartan-treated adult SHR (**Figs. 5** and **6; Suppl Fig. S2**). The number of PRV-labeled neurons in the spinal IML did not differ significantly between vehicle-treated and losartan-treated SHR **(Suppl Fig. S2)**. Strikingly, the numbers of PRV-labeled neurons in both the RVLM and PVN were substantially reduced in losartan-treated SHR compared with vehicle-treated SHR (n = 6 rats per group, RVLM: *P* = 0.00096, *t*_(10)_ = 4.61; PVN: *P* = 0.0011, *t*_(10)_ = 4.51; **Figs. 5** and **6**).

**Fig. 6.**
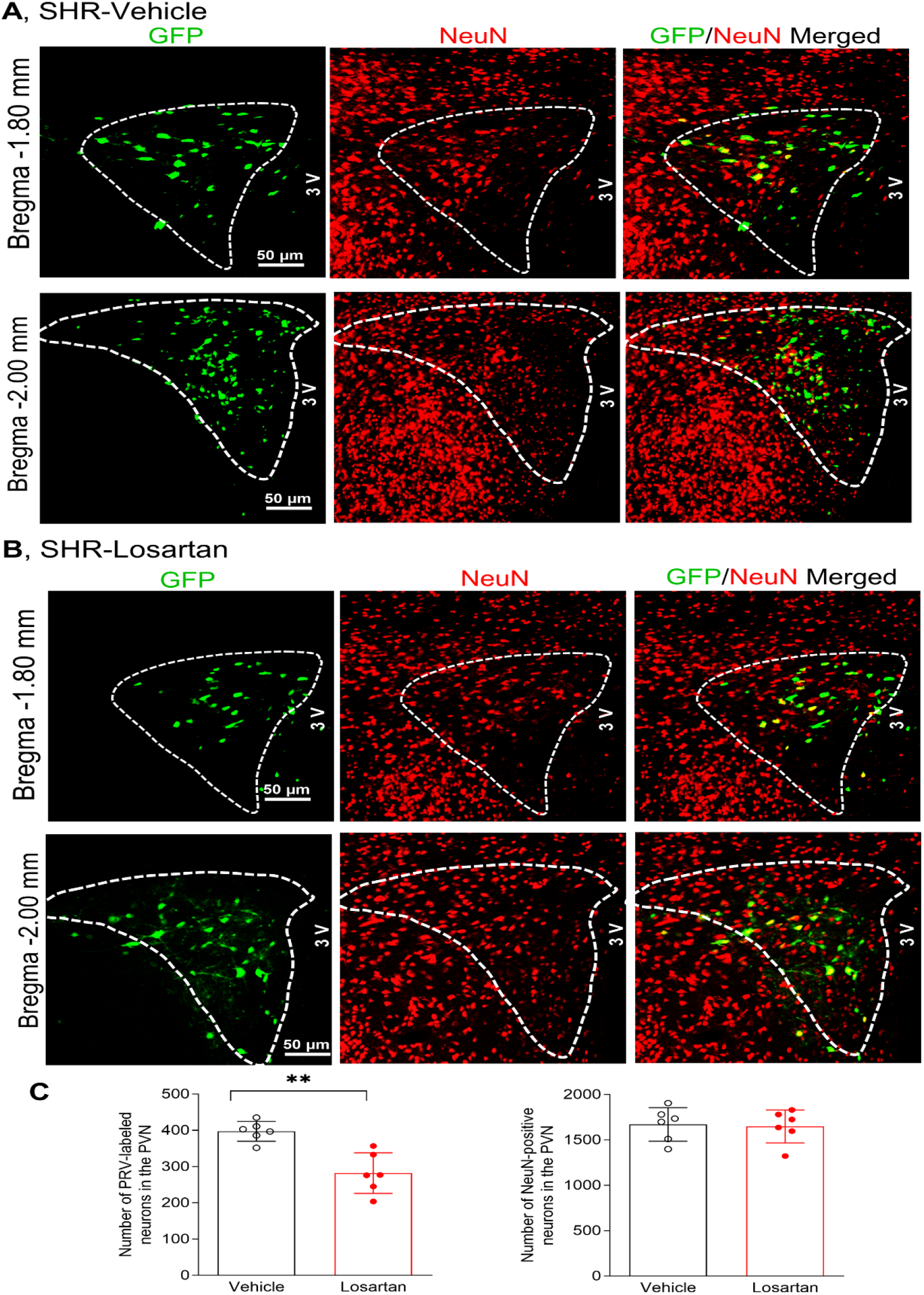
Long-term treatment with losartan reduces the increased number of presympathetic neurons innervating the adrenal gland in the PVN of SHR. **A** and **B**, Representative confocal images show presympathetic neurons (dual-labeled by GFP and NeuN) in the PVN of vehicle- or losartan-treated SHR (50 mg/kg per day) from 4 to 12 weeks of age followed by PRV-injection into the left adrenal gland. Scale bar, 50 µm. **C**, Summary data show the number of presympathetic neurons and NeuN-immunoreactive neurons in the PVN in SHR treated with vehicle or losartan (n = 6 rats per group; \*\**P* < 0.01, two-tailed Student’s *t* test).

In contrast, both the distribution and numbers of PRV-labeled neurons in the spinal IML, RVLM, and PVN were comparable between vehicle-treated and losartan-treated adult WKY (n = 6 rats per group, **Suppl Figs. S2-S4**). Together, these results suggest that heightened AT1 receptor activity during the development of hypertension contributes to the expansion of brain presympathetic neural networks in SHR.

### Long-term treatment with losartan in young SHR blocks the elevated active glutamatergic synaptic contacts onto PVN presympathetic neurons

Finally, we investigated the mechanisms by which increased AT1 receptor activity promotes the expansion of presympathetic neural networks in the brains of SHR. The increase in presympathetic neurons projecting to the adrenal gland, without a change in total neuronal number, suggests that sustained AT1 receptor activation enhances presympathetic synaptic connectivity in the PVN and RVLM likely through synaptogenesis. Because fluorescence-labeled neurons can be visualized in the PVN, but not in the RVLM or spinal IML, in live tissue slices from adult rats, we examined glutamatergic synaptic activity of presympathetic neurons in the PVN and assessed the effect of long-term losartan treatment. Losartan (50 mg/kg per day in drinking water) or vehicle was administered to SHR and WKY rats from 4 to 12 weeks of age. PVN presympathetic neurons were retrogradely labeled by injecting FluoSpheres, a fluorescent tracer (27, 54), into the spinal IML of 12-week-old of SHR and WKY treated with losartan or vehicle. Because PRV can be cytotoxic to labeled neurons, we used a non-viral fluorescence tracer to identify presympathetic neurons in the PVN for brain slice recordings. We recorded miniature excitatory postsynaptic currents (mEPSCs), which represent spontaneous, quantal release of glutamate from presynaptic terminals, in fluorescence-labeled PVN neurons. An increase in mEPSC frequency, without a change in amplitude, can reflect an increase in the number of active excitatory synapses (55). Consistent with previous reports (26, 43, 56), vehicle-treated SHR exhibited a higher baseline frequency, but not amplitude, of mEPSCs in labeled PVN neurons compared with vehicle-treated WKY (n = 12 neurons per group, *F*_(3,44)_ = 30.45, *P* < 0.001; **Fig. 7A-C**). Remarkably, long-term losartan treatment largely normalized the increased mEPSC frequency in labeled PVN neurons of adult SHR (n = 12 neurons per group, *F*_(3,44)_ = 30.45, *P* < 0.001; **Fig. 7A-C**). In contrast, the baseline frequency and amplitude of mEPSCs of labeled PVN neurons did not differ significantly between vehicle-treated and losartan-treated WKY (n = 12 neurons per group, **Fig. 7A-C**). These findings collectively suggest that enhanced AT1 receptor activity during hypertension development contributes to the increased number of active glutamatergic synaptic contacts onto PVN presympathetic neurons in SHR.

**Fig. 7.**
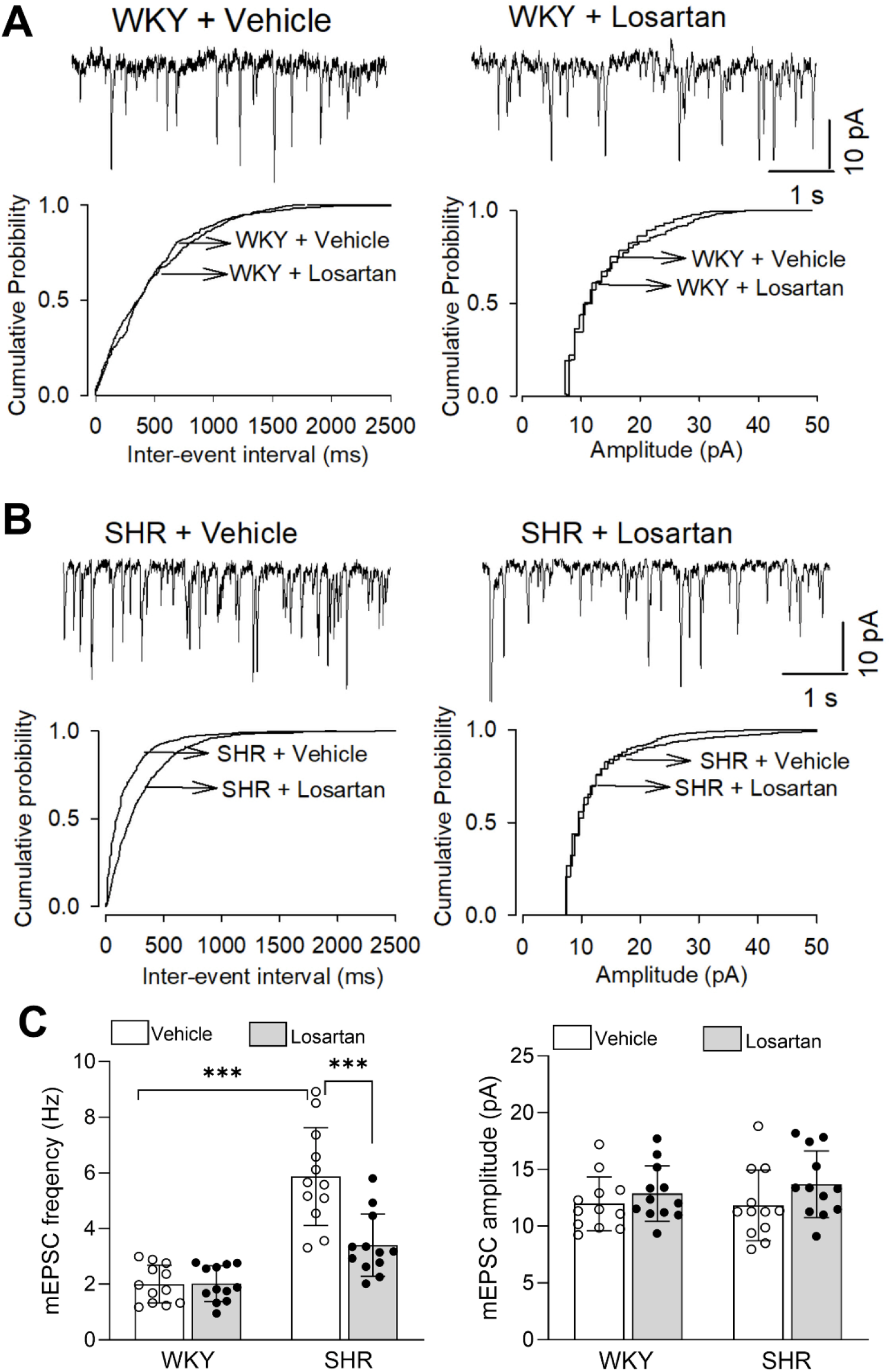
Long-term treatment with losartan normalizes the elevated active glutamatergic synaptic contacts onto PVN presympathetic neurons in SHR. **A** and **B**, Representative mEPSC recording traces and corresponding cumulative plots of the frequency and amplitude of mEPSCs in spinally projecting PVN presympathetic neurons of WKY (**A**) and SHR (**B**) treated with vehicle or losartan (50 mg/kg per day) from 4 to 12 weeks of age. **C**, Summary data show the mean frequency and amplitude of mEPSCs in spinally projecting PVN neurons from WKY and SHR treated with vehicle or losartan (n = 12 neurons from 3 rats per group; \*\*\**P* < 0.001, two-way ANOVA with Tukey’s *post hoc* test).

## Discussion

Our study demonstrates, for the first time, that presympathetic neuronal connectivity in the RVLM and PVN is expanded in SHR, a genetic model of essential hypertension. PRV-152, a genetically engineered virus expressing GFP, is a unique tool for mapping neural circuits as it specifically infects neurons and spreads retrogradely across synapses (21, 37). Because PRV can transverse multiple synapses, it reveals chains of neurons connected to the initial injection site. PRV does not infect humans or other primates, making it safer for laboratory use. Although presympathetic neural circuits have been well mapped in the normotensive rat brain using PRV (21-23), whether these networks are altered under hypertensive conditions remains unknown. In this study, we selected the adrenal gland for injection of PRV-152 to assess presympathetic neuronal connectivity in SHR, because it is a sympathetically innervating end organ critically involved in enhanced sympatho-adrenal activation and blood pressure elevation (57-59). Functionally, the adrenal medulla acts as an extension of the sympathetic nervous system. It receives direct input from spinal preganglionic neurons, bypassing peripheral ganglia, and releases epinephrine and norepinephrine into the bloodstream, producing a widespread sympathetic response. The PVN coordinates sympathetic output through projections to the RVLM and IML in the spinal cord (7, 12, 22). We found that the numbers of PRV-labeled neurons in both the RVLM and PVN were substantially higher in adult SHR than in normotensive controls. This expanded presympathetic connectivity is associated with enhanced sympatho-adrenal activation in SHR (44, 58). Interestingly, this expansion was absent in young, prehypertensive SHR. Because hypertension development in SHR is age-dependent, the age-related expansion of presympathetic neuronal connectivity likely contributes to increased sympathetic vasomotor activity and the development of hypertension.

A striking finding of our study is that potentiated AT1 receptor activity in SHR plays a crucial role in this expansion of presympathetic neuronal connectivity in the PVN and RVLM during hypertension development. Sustained blockade of AT1 receptors from a young age largely prevents the development of hypertension in SHR (17, 29). Angiotensin and AT1 receptors are abundantly expressed in forebrain regions, particularly the PVN (60, 61). While AT1 receptor expression in the PVN is comparable between young, prehypertensive SHR and WKY, it increases markedly in adult SHR (3). Circulating angiotensin II may also cross a compromised blood-brain barrier, further enhancing AT1 receptor activity in the brain of adult SHR (62). The age-dependent elevation in AT1 receptor expression and activity in the PVN and RVLM contributes to augmented sympathetic outflow and the progression and maintenance of hypertension in SHR (3, 17, 52). In this study, we showed that presympathetic neuronal connectivity in the brain was unchanged in young SHR but expanded in adult SHR. This temporal correlation between increased AT1 receptor expression/activity and expanded presympathetic neuronal connectivity suggests a potential role for AT1 receptors in regulating presympathetic circuitry in the brain. Supporting this notion, we found that prolonged blocking of AT1 receptors with losartan in SHR, initiated during the prehypertensive stage, diminished the development of hypertension and concurrently reversed the increase in PRV-labeled neurons in the PVN and RVLM in adult SHR. Notably, losartan treatment had no noticeable effect on the number of PRV-labeled neurons in these brain regions in normotensive control rats. Because losartan crosses the blood-brain barrier (25, 32), our findings suggest that potentiated local RAS and AT1 receptor activity promote the age-dependent expansion of presympathetic neural circuits, thereby contributing to hypertension development in SHR.

Our study provides new functional evidence that overactive AT1 receptors enhance excitatory glutamatergic synaptic activity onto PVN presympathetic neurons in SHR. Because the total number of neurons in the PVN and RVLM was comparable between young and adult SHR, as well as between adult WKY and adult SHR, the increased number of PRV-labeled neurons observed in adult SHR is likely attributable to the recruitment of additional presympathetic neurons in the PVN and RVLM via enhanced synaptogenesis. Consistent with this interpretation, excitatory glutamatergic synaptic input to PVN presympathetic neurons is increased in adult SHR and contributes to excessive sympathetic outflow (13, 43, 56). We focused our electrophysiological recordings on presympathetic neurons in the PVN because the spinal IML and RVLM of adult rats are heavily myelinated, precluding visualization of fluorescence-labeled neurons in live spinal and brainstem slices. Although PRV is an attenuated viral tracer, it retains cytotoxicity, particularly with prolonged post-infection survival (37). To circumvent this limitation, we used a non-viral fluorescence tracer to retrogradely label PVN presympathetic neurons for brain slice recordings. Using this approach, we found that the frequency—but not the amplitude—of mEPSCs in PVN presympathetic neurons was substantially increased in adult SHR. Remarkably, long-term treatment with losartan largely normalized this increased mEPSC frequency of PVN presympathetic neurons in SHR but had no effect in WKY. mEPSCs are widely used to assess synaptic strength and functional connectivity of neurons. mEPSCs provide a well-established measure of synaptic strength and functional connectivity. In general, changes in mEPSC frequency reflect alterations in the number of functional synapses or presynaptic release probability, whereas changes in amplitude primarily indicate postsynaptic responsiveness, often associated with altered AMPA receptor expression. Thus, our findings indicate that enhanced AT1 receptor signaling increases the number of active excitatory glutamatergic synapses onto PVN presympathetic neurons in SHR.

The exact signaling mechanisms linking AT1 receptor activation to glutamatergic synaptogenesis in SHR remain incompletely defined. Increased glutamate NMDA receptor activity is likely a key contributor, because enhanced synaptic activity promotes input-specific dendritic remodeling and long-term stabilization of synapses through NMDA receptors (63, 64). In cortical neurons, angiotensin II enhances NMDA receptor activity through AT1 receptors (65). In the PVN and RVLM of adult SHR, the activities of both AT1 receptors and NMDA receptors are elevated (3, 43, 52, 66). Synaptic NMDA receptor activity in PVN presympathetic neurons is further augmented reciprocally by increased protein kinase activity and/or diminished calcineurin activity in SHR (67-70). Additionally, potentiated AT1 receptor activity promotes synaptic localization and activity of α2δ-1–dependent NMDA receptors through PKC-mediated phosphorylation in these neurons in SHR (26). Together, these findings support a model in which age-dependent increases in AT1 receptor expression and activity drive NMDA receptor–mediated glutamatergic synaptogenesis in PVN presympathetic neurons during the development of hypertension in SHR. Sustained AT1 receptor blockade may therefore attenuate hypertension progression by limiting NMDA receptor– dependent synaptic connectivity within presympathetic circuits. Enhanced glutamatergic synaptic drive, combined with expansion of presympathetic circuitry, may stabilize and amplify sympathetic output, thereby contributing to chronic sympathetic overactivation in hypertension.

AT1 receptors are widely expressed throughout the brain and peripheral tissues, including the kidneys and blood vessels (71). By binding to the AT1 receptor, angiotensin II can cause vasoconstriction and enhance salt and water retention via aldosterone secretion from the adrenal gland (72-74). Our findings suggest that the reduced expansion of presympathetic neuronal connectivity and diminished glutamatergic synaptic activity in presympathetic neurons of SHR by long-term losartan treatment are likely mediated, at least in part, by inhibition of brain AT1 receptor activity. Consistent with interpretation, systemically administered losartan blocks AT1 receptor binding and activity within the brain (31, 32). Furthermore, increasing DNA methylation in the PVN of SHR markedly lowers arterial blood pressure and reduces AT1 receptor expression (3), indicating that the age-dependent rise in AT1 receptor expression is driven by DNA demethylation rather than elevated arterial pressure per se. In line with this conclusion, lowering arterial pressure via celiac ganglionectomy does not reduce the elevated mEPSC frequency in PVN presympathetic neurons of SHR (69), suggesting that enhanced glutamatergic synaptic input to these neurons is unlikely to be a secondary consequence of hypertension. Nevertheless, because reduced sympathetic outflow to the kidney and adrenal gland decreases RAS activity by inhibiting renin release from the kidneys and suppressing the system’s overall response (4), the antihypertensive effects of losartan are likely mediated by its reciprocal inhibitory actions on both central and peripheral AT1 receptors.

In summary, our study provides novel anatomical (transsynaptic labeling) and physiological (synaptic activity recordings) evidence for a maladaptive, angiotensin-dependent expansion and reinforcement of brain presympathetic pathways in SHR. Enhanced AT1 receptor signaling drives both structural enlargement and excitatory synaptic strengthening of presympathetic networks in the PVN and RVLM. This pathological remodeling of presympathetic circuitry likely underlies the heightened sympathetic vasomotor tone characteristic of neurogenic hypertension. Moreover, these findings provide mechanistic insight into the antihypertensive efficacy of long-term AT1 receptor antagonism, highlighting its potential to reduce sympathetic outflow by reversing hyperactive presympathetic networks in the brain.

## Acknowledgements

We are grateful to Dr. Lynn Enquist at Princeton University for providing PRV-152 used in this study. This work was supported by a grant (R01HL154512) from the National Institutes of Health and by the Pamela and Wayne Garrison Distinguished Chair Endowment.

## Supplemental Figures

**Suppl Fig. S1.**
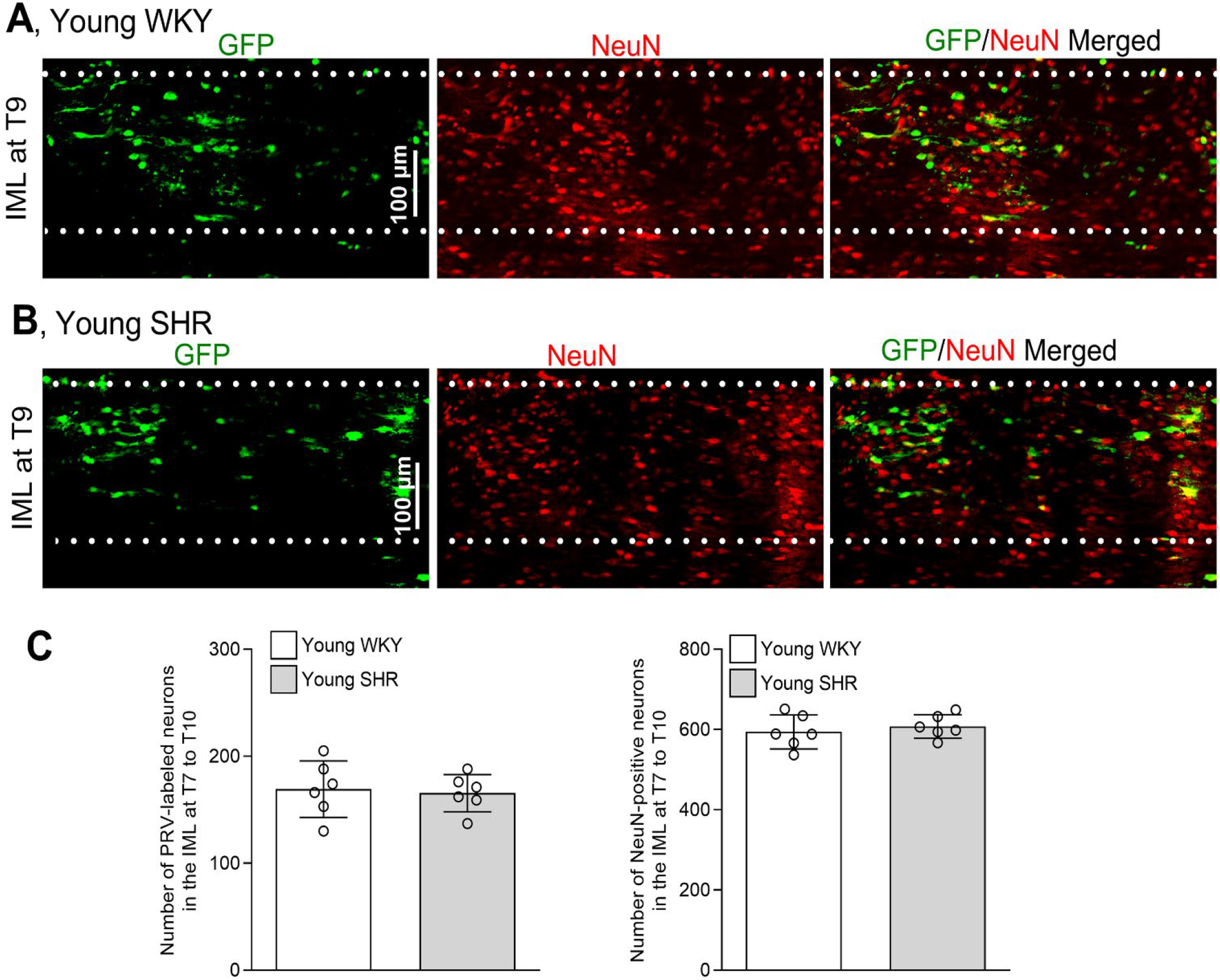
The number of preganglionic neurons in the spinal IML that innervate the adrenal gland is similar between young SHR and WKY. **A** and **B**: Representative confocal images of sagittal spinal cord sections at the T9 level show preganglionic neurons (dual-labeled by GFP and NeuN) in the spinal IML in young WKY (**A**) and SHR (**B**) with PRV-152 injected into the left adrenal gland. Scale bar, 100 µm. **C**, Summary data show the numbers of preganglionic neurons (dual-labeled by GFP and NeuN) and NeuN immunoreactive neurons in the spinal IML from T7 to T10 in young WKY and SHR (n = 6 rats per group; two-tailed Student’s *t* test).

**Suppl Fig. S2.**
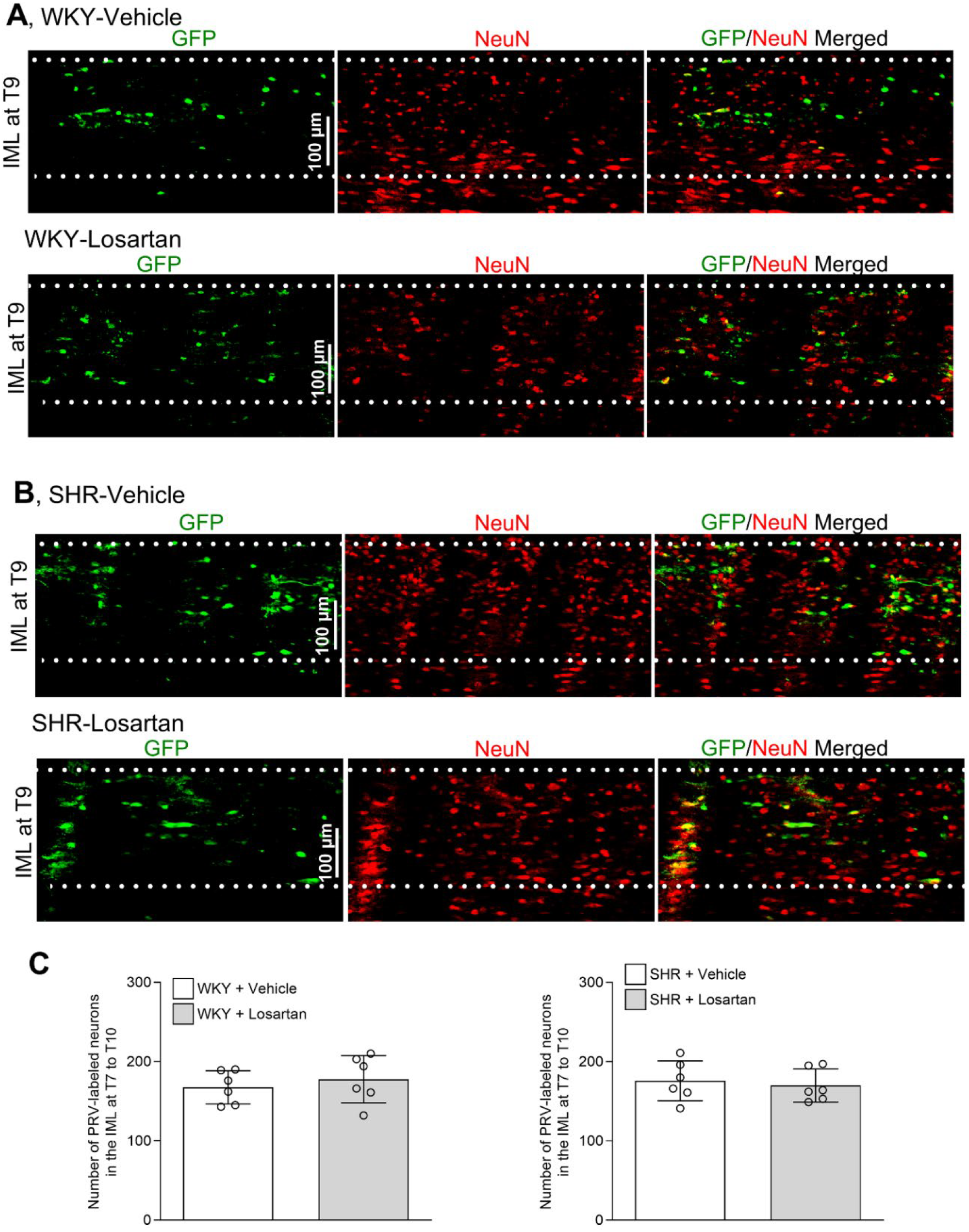
Long-term treatment with losartan does not affect the number of preganglionic neurons in the spinal IML of SHR and WKY. **A** and **B**, Representative confocal images of sagittal spinal cord sections at the T9 level show preganglionic neurons (dual-labeled by GFP and NeuN) in the spinal IML in vehicle- or losartan-treated WKY (**A**) and SHR (**B**) with PRV-152 injected into the left adrenal gland. Scale bar, 100 µm. **C**, Summary data show the mean numbers of preganglionic neurons (dual-labeled by GFP and NeuN) in the spinal IML from T7 to T10 in vehicle- or losartan-treated WKY and SHR with PRV-152 injected into the left adrenal gland (n = 6 rats per group; two-tailed Student’s *t* test).

**Suppl Fig. S3.**
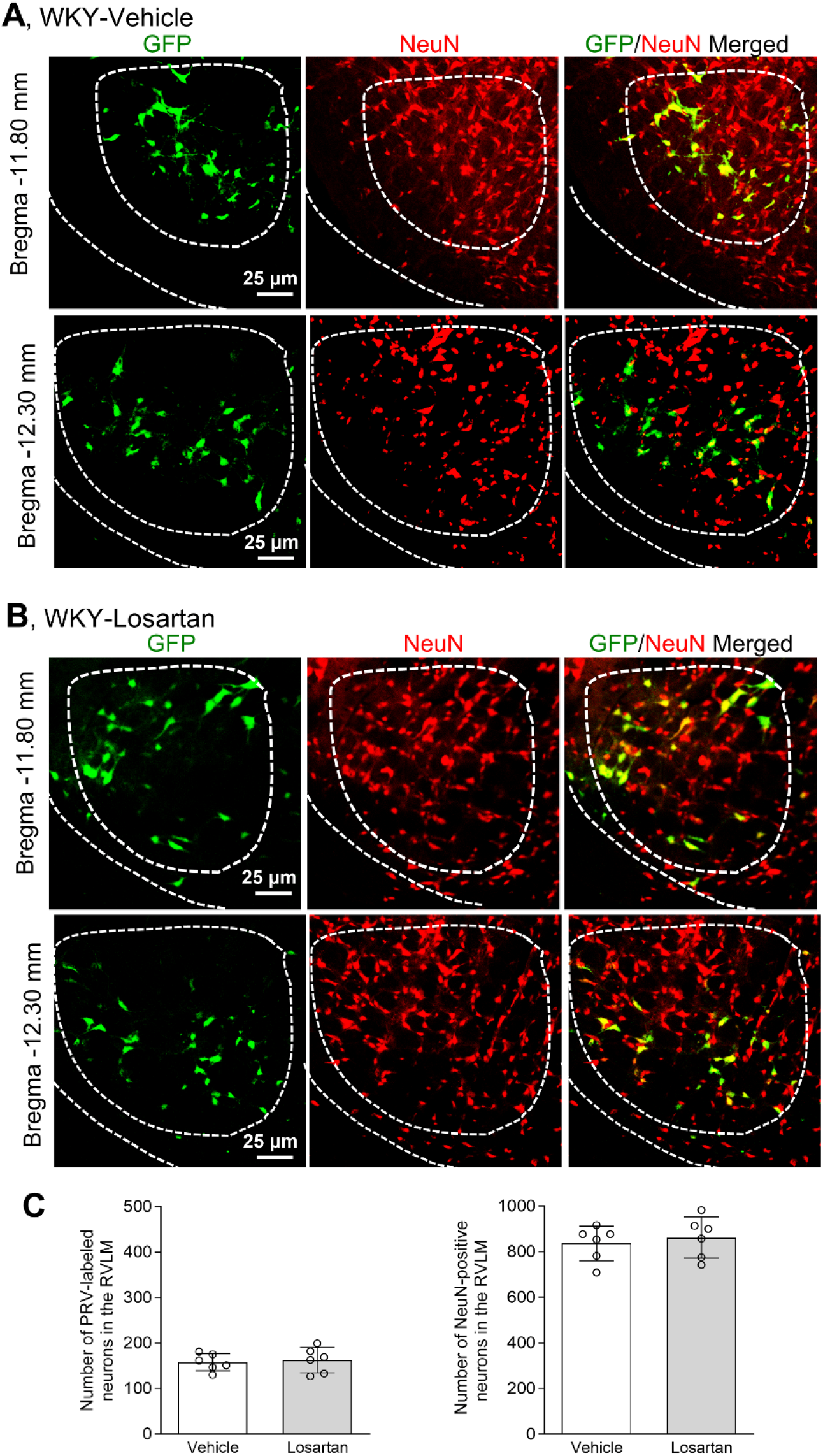
Prolonged treatment with losartan does not affect the number of presympathetic neurons innervating the adrenal gland in the RVLM of WKY. **A** and **B**, Representative confocal images show presympathetic neurons (dual-labeled by GFP and NeuN) in the RVLM of WKY treated with vehicle or losartan from 4 to 12 weeks of age followed by PRV-152 injected into the left adrenal gland. Scale bar, 25 µm. **C**, Summary data show the number of presympathetic neurons (dual-labeled by GFP and NeuN) and NeuN immunoreactive neurons in the RVLM of vehicle- or losartan-treated WKY with PRV-152 injected into the left adrenal gland (n = 6 rats per group, two-tailed Student’s *t* test).

**Suppl Fig. S4.**
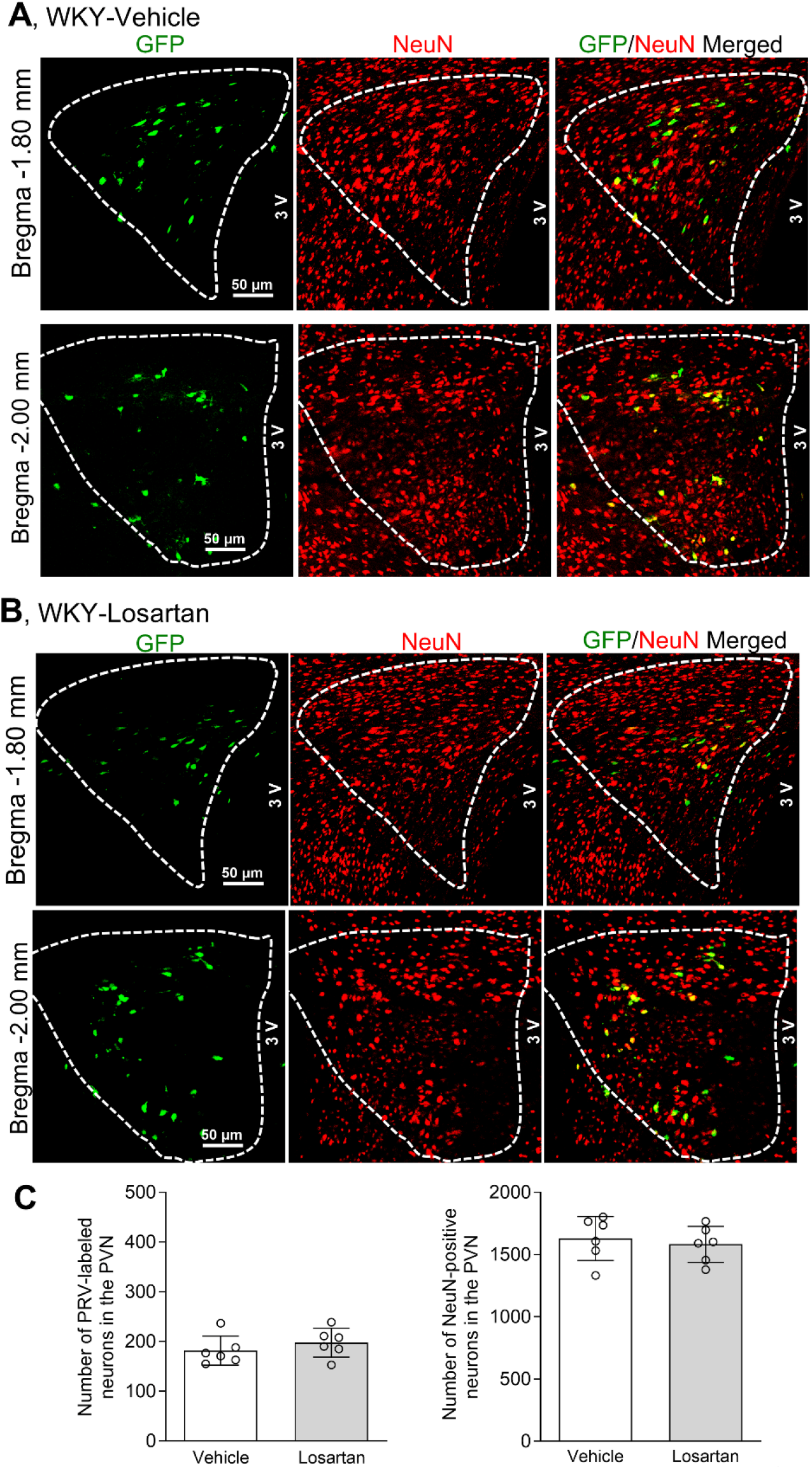
Long-term treatment with losartan does not affect the number of presympathetic neurons innervating the adrenal gland in the PVN of WKY. **A** and **B**, Representative confocal images show presympathetic neurons (dual-labeled by GFP and NeuN) in the PVN of WKY treated with vehicle or losartan from 4 to 12 weeks of age followed by PRV-152 injected into the left adrenal gland. Scale bar, 50 µm. **C**, Summary data show the number of presympathetic neurons (dual-labeled by GFP and NeuN) and NeuN immunoreactive neurons in the PVN of vehicle- or losartan-treated WKY with PRV-152 injected into the left adrenal gland (n = 6 rats per group, two-tailed Student’s *t* test).

